# Urinary protein changes during the short-term growth and development of rats

**DOI:** 10.1101/2023.05.25.542266

**Authors:** Yuqing Liu, Minhui Yang, Haitong Wang, Yuzhen Chen, Youhe Gao

## Abstract

Can the urine proteome reflect short-term changes in the growth and development of animals? Do short-term developmental effects on urinary protein need to be considered when performing urine marker studies using model animals with faster growing periods? In this study, urine samples were collected from 10 Wistar rats aged 6-8 weeks 3 and 6 days apart. The results showed that the urine proteome could sensitively reflect short-term growth and development in rats. For example, comparing the urine proteome of Day 0 and Day 6, 195 differential proteins were identified after screening (FC ≥ 1.5 or ≤ 0.67, P < 0.05), and verified by randomization, the average number of randomly generated differential proteins was 17.99. At least 90.77% of the differential proteins were not randomly generated. This finding demonstrates that the differential proteins identified in the samples collected at different time points were not randomly generated. A large number of biological processes and pathways related to growth and development were enriched, which shows that the urine proteome reflects the short-term growth and development of rats, and provides a means for in-depth and meticulous study of growth and development. Moreover, an interfering factor in animal experiments using 6-to 8-week-old rats to construct models was identified. The results of this study demonstrated that there were differences in the urinary proteome in rats aged 6-8 weeks only 3-6 days apart, which suggests that the sensitivity of urinary proteomics is high and shows the sensitive and precise response of the urinary proteome to body changes.

## 1. Introduction

### 1.1 Effects of Growth and Development on the Urine Proteome in Rats

To date, few studies have used urine proteomics to explore the growth and development of rats. In 2023, Pan Xuanzhen et al.^[1]^ first used urinary proteomic technology to track the changes at several important developmental time points in a group of rats, covering 10 time points from childhood, adolescence, young adulthood, mid-adulthood, old age, and close to death, providing new information, filling the gap of the field of developmental research. They found changes in several organs and aging-related pathways through all of the time pointes, demonstrated for the first time that urine can reflect changes in various aspects of body growth and development, and provided ideas for monitoring the patient’s physical condition, including clinical prognosis, and future aging studies^[1]^.

### 1.2 Urine Biomarkers

Biomarkers are indicators that can objectively reflect normal pathological processes as well as physiological processes^[2]^, and clinically, biomarkers can predict, monitor, and diagnose multifactorial diseases at different stages^[3]^. The potential of urinary biomarkers has not been fully developed compared to more widely used blood biomarkers. However, urinary biomarkers are especially important in terms of early diagnosis of disease and prediction of status. Since homeostatic mechanisms are regulated in the blood, changes in the blood proteome caused by disease are metabolically excreted, and no significant changes can be apparent in the early stages of the disease. Whereas urine is produced by glomerular filtration of plasma and is not regulated by homeostatic mechanisms, and minor changes in the disease at an early stage can be observed in urine. According to a previous study, the changes observed in urine occur far earlier than those in blood, earlier than those in pathological sections, and even earlier than the appearance of disease symptoms, suggesting that urine biomarkers can be applied for the early diagnosis of diseases. Since the urine proteome is susceptible to a variety of factors, such as diet, drug therapy, and daily activities, to make the experimental results more accurate, the key is to use a simple and controllable system. The genetic and environmental factors of animal models can be artificially controlled and can minimize the influence of unrelated factors, indicating that animal models are a very appropriate experimental method. For example, (1) Zhang Fansheng et al.^[4]^ found that the levels of 29 proteins changed in the urine before amyloid plaque deposition in the brains of transgenic mice with Alzheimer’s disease appeared, and 24 of them were reported to be associated with Alzheimer’s disease or used as markers. (2) Wu Jianqiang et al.^[5]^ demonstrated that in a Walker 256 subcutaneous tumor rat model, the levels of 10 proteins changed in the urine before the subcutaneous tumor was palpable. (3) Zhang Y et al.^[6]^ illustrated that in a chronic pancreatitis rat model, 15 differential proteins were identified in the urine before the pathology had changed at week 2, of which 5 were reported to be associated with pancreatitis; (4) Ni Yanying et al.^[7]^ showed that in a glioma rat model constructed by C6 cell injection into the brain, urinary protein levels changed before the appearance of pathological changes during imaging; (5) Zhang Fansheng et al.^[8]^ found that in a thioacetamide-induced liver fibrosis rat model, 40 differential proteins were identified in the urine before pathological changes, and 15 were reported to be associated with fibrosis. (6) Yin et al.^[9]^ found that urine glucose levels presented frequent disturbances before blood glucose increased in obese type 2 diabetic rats, which is important in indicating early diabetes. (7) Huang He et al.^[10]^ exposed rats to smoke from traditional cigarettes and screened biomarkers of chronic obstructive pulmonary disease (COPD) that have been reported only at two weeks of exposure. After comparative studies, it was found that urinary protein levels changed when tumor cells were subcutaneously injected^[5]^ and grew in different organs including liver^[8]^, bone^[11]^, lung^[12]^, and brain^[7]^, which indicated that urine had the potential to distinguish the growth of the same tumor cells in different organs. In terms of sample acquisition, urine acquisition is more noninvasive and easily available^[13]^, urine is a suitable source of biomarkers, and the construction of animal models is a very important means in urine proteomics research.

However, the growth and development of animals during the experiment is an important influencing factor that cannot be ignored. Therefore, can the urinary proteome reflect changes in the short-term growth and development of rats? Is it necessary to consider the effect of short-term development on urinary proteins when performing marker studies using model animals that grow more rapidly? This is a challenge to the sensitivity boundary of urinary proteomics. Therefore, 6-to 8-week-old Wistar rats with rapid growth and development were selected to study the dynamic changes in growth and development status reflected in the urine proteome within one week (Figure 1). This study investigated the degree to which the urine proteome precisely reflects growth and development. The effect of growth and development, which has been neglected, on animal model experiments was explored.

**Figure 1.**
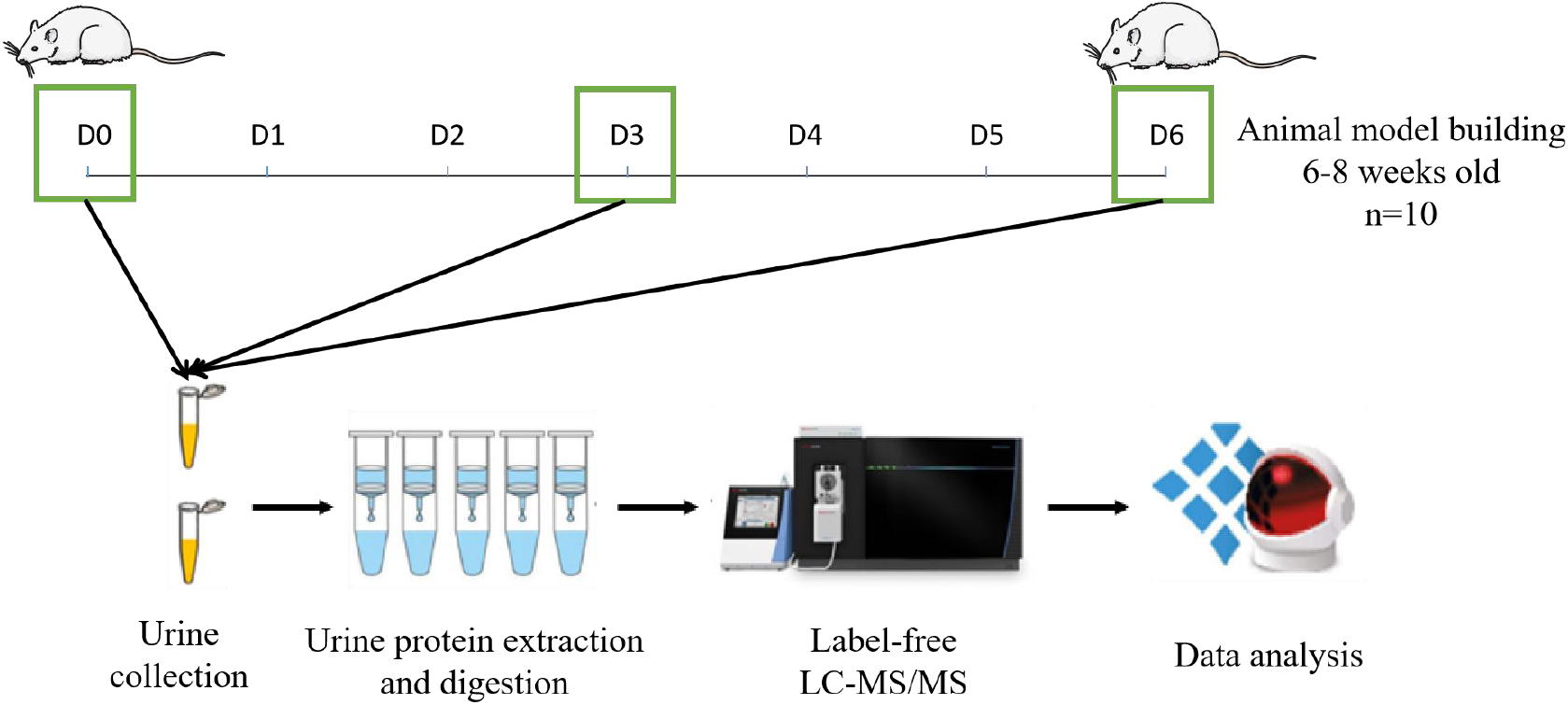
Workflow for urine proteomics analysis during the short-term growth and development of rats. Urine samples were collected on Days 0, 3, and 6. After urine samples were collected and processed, the protein groups were identified using liquid chromatography coupled with tandem mass spectrometry (LC–MS/MS) to quantitatively analyze changes caused by short-term growth and development in rats.

## 2. Materials and Methods

### 2.1 Urine collection

In this experiment, 10 healthy male Wistar rats (180-200 g) of SPF grade aged 6-8 weeks were purchased from Beijing Vital River Laboratory Animal Technology Co., Ltd., with an animal license number of SYXK (jing) 2021-0011. All rats were maintained in a standard environment (room temperature (22 ± 1) °C, humidity 65%–70%). All rats were kept in a new environment for three days before starting the experiment, and all experimental procedures followed the review and approval of the Ethics Committee of the College of Life Sciences, Beijing Normal University, with approval number CLS-AWEC-B-2022-003. Rats were observed for behavioral changes during the experiment, and body weights were recorded every 3 days.

After all rats were kept in a new environment for three days, they were uniformly placed in metabolic cages to collect urine samples for 12 h, i.e., Day 0 samples. All rats were placed in metabolic cages on Days 3 and 6 to collect urine samples for 12 h. Rats were fasted and water deprived during urine collection, and all collected urine samples were stored in a -80 °C freezer.

### 2.2 Treatment of the urine samples

Urine protein extraction and quantification: Rat urine samples collected at three time points were centrifuged at 12,000×g for 40 min at 4 °C, and the supernatants were transferred to new Eppendorf (EP) tubes. Three volumes of precooled absolute ethanol were added, homogeneously mixed and precipitated overnight at -20 °C. The following day, the mixture was centrifuged at 12,000×g for 30 min at 4 °C, and the supernatant was discarded. The protein pellet was resuspended in lysis solution (containing 8 mol/L urea, 2 mol/L thiourea, 25 mmol/L dithiothreitol, and 50 mmol/L Tris). The samples were centrifuged at 12,000×g for 30 min at 4 °C, and the supernatant was placed in a new EP tube. The protein concentration was measured using the Bradford assay.

Urinary protease cleavage: A 100 μg urine protein sample was added to the filter membrane (Pall, Port Washington, NY, USA) of a 10 kDa ultrafiltration tube and placed in an EP tube, and 25 mmol/L NH4HCO3 solution was added to make a total volume of 200 μL. Then, 20 mM dithiothreitol solution (dithiothreitol, DTT, Sigma) was added, and after vortex mixing, the metal bath was heated at 97 °C for 5 min and cooled to room temperature. Iodoacetamide (IAA, Sigma) was added at 50 mM, mixed well and allowed to react for 40 min at room temperature in the dark. Then, membrane washing was performed: ① 200 μL of UA solution (8 mol/L urea, 0.1 mol/L Tris-HCl, pH 8.5) was added and centrifuged twice at 14,000×g for 5 min at 18 °C; ② Loading: freshly treated samples were added and centrifuged at 14,000×g for 40 min at 18 °C; ③ 200 μL of UA solution was added and centrifuged at 14,000×g for 40 min at 18 °C, repeated twice; ④ 25 mmol/L NH4HCO3 solution was added and centrifuged at 14,000×g for 40 min at 18 °C, repeated twice; ⑤ trypsin (Trypsin Gold, Promega, Trypchburg, WI, USA) was added at a ratio of 1:50 trypsin:protein for digestion and kept in a water bath overnight at 37 °C. The following day, peptides were collected by centrifugation at 13,000×g for 30 min at 4 °C, desalted through an HLB column (Waters, Milford, MA), dried using a vacuum dryer, and stored at -80 °C.

### 2.3 LC-MS/MS analysis

The digested samples were reconstituted with 0.1% formic acid, and the peptide sample concentrations were quantified using a BCA kit. They were then diluted to 0.5 μg/μL. Mixed peptide samples were prepared from 4 μL of each sample and separated using a high pH reversed-phase peptide separation kit (Thermo Fisher Scientific) according to the instructions. Ten effluents (fractions) were collected by centrifugation, dried using a vacuum dryer and reconstituted with 0.1% formic acid. iRT reagent (Biognosys, Switzerland) was added at a volume ratio of sample:iRT of 10:1 to calibrate the retention times of extracted peptide peaks. For analysis, 1 μg of each peptide from an individual sample was loaded onto a trap column and separated on a reverse-phase C18 column (50 μm×150 mm, 2 μm) using the EASY-nLC1200 HPLC system (Thermo Fisher Scientific, Waltham, MA). The elution for the analytical column lasted 90 min with a gradient of 5%-28% buffer B (0.1% formic acid in 80% acetonitrile; flow rate 0.3 μL/min). Peptides were analyzed with an Orbitrap Fusion Lumos Tribrid Mass Spectrometer (Thermo Fisher Scientific, MA).

To generate the spectrum library, 10 isolated fractions were subjected to mass spectrometry in data-dependent acquisition (DDA) mode. Mass spectrometry data were collected in high sensitivity mode. A complete mass spectrometric scan was obtained in the 350-1500 m/z range with a resolution set at 60,000. Individual samples were analyzed using Data Independent Acquisition (DIA) mode. DIA acquisition was performed using a DIA method with 36 windows. After every 10 samples, a single DIA analysis of the pooled peptides was performed as a quality control.

### 2.4 Database searching and label-free quantitation

Raw data collected from liquid chromatography-mass spectrometry were imported into Proteome Discoverer (version 2.1, Thermo Scientific) and the Swiss-Prot rat database (published in May 2019, containing 8086 sequences) for alignment, and iRT sequences were added to the rat database. Then, the search results were imported into Spectronaut Pulsar (Biognosys AG, Switzerland) for processing and analysis. Peptide abundances were calculated by summing the peak areas of the respective fragment ions in MS^2^. Protein intensities were summed from their respective peptide abundances to calculate total protein abundances.

### 2.5 Statistical analysis

Two technical replicates were performed for each sample, and the average was used for statistical analysis. In this experiment, samples at different time periods were contrasted. Identified proteins were contrasted to screen for differential proteins. There were two differential protein screening conditions. Loose conditions were as follows: fold change between groups (FC, fold change) ≥ 1.5 or ≤ 0.67, P value < 0.05 by two-tailed unpaired t test analysis. Strict conditions were as follows: FC ≥ 2 or ≤ 0.5, P value < 0.01. Randomization was also used to test the reliability of the selected differential proteins to demonstrate that the selected differential proteins between groups were indeed produced by the growth and development of the rats themselves rather than produced randomly. The Wukong platform was used for the selected differential proteins (https://www.omicsolution.com/wkomics/main/), and the UniProt website (Release 2023_01) (https://www.uniprot.org/) and the OmicsBean database (http://www.omicsbean.cn/) were used to perform data processing. The biological function enrichment analysis was conducted using STRING (https://cn.string-db.org/), a database that performs the analysis of protein interaction networks. In the PubMed database (https://pubmed.ncbi.nlm.nih.gov), the reported literature was searched to conduct the functional analysis of the differential proteins.

## 3. Results

### 3.1 Growth and development characteristics in rats

In this experiment, body weight was monitored during the growth and development of rats, and the body weight of rats was recorded every 3 days (Figure 2). Morever, the rats were lively and active, with normal dietary water intake and no abnormal findings.

**Figure 2.**
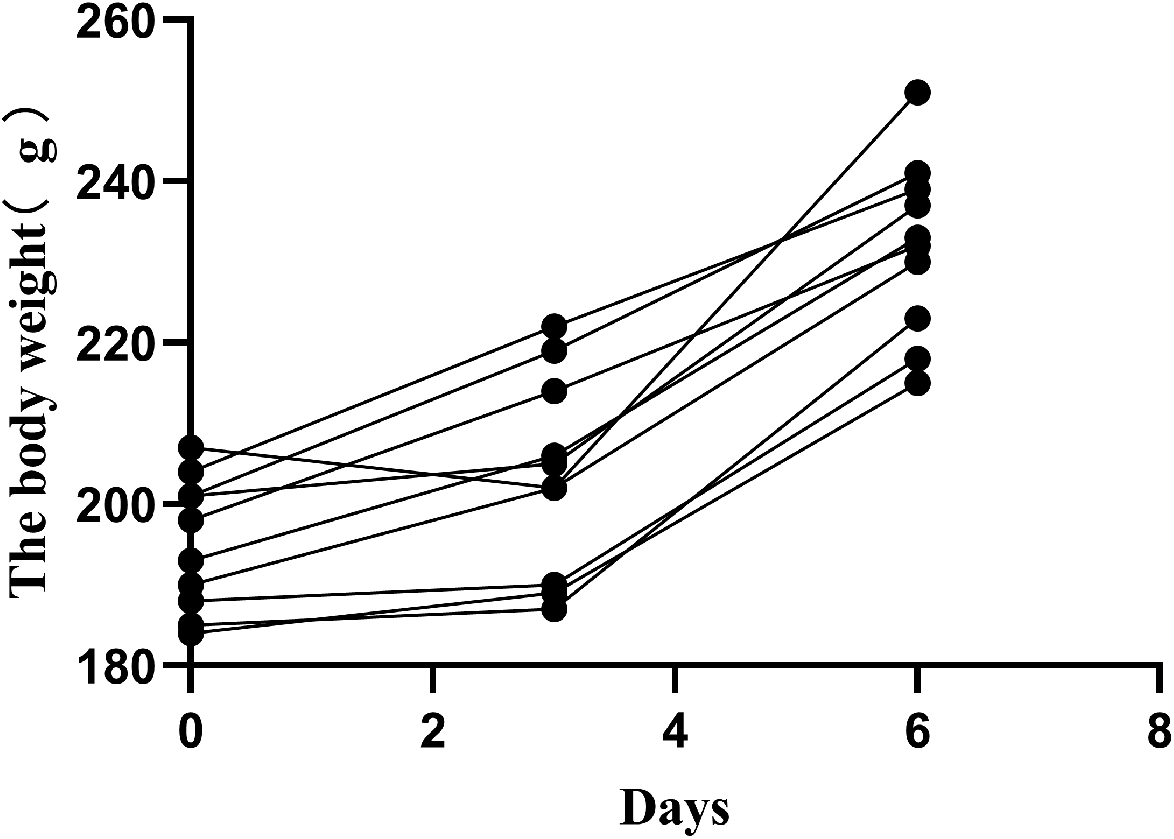
Short-term growth and developmental weight changes in 10 male Wistar rats aged 6-8 weeks.

### 3.2 Analysis of urinary proteome changes during the short-term growth and development of rats

#### 3.2.1 Urine protein identification

LC-MS/MS tandem mass spectrometry was performed on peptides resulting from the digestion of 30 urine samples collected from 10 Wistar male rats 6-8 weeks of age following urine sample collection on Days 0, 3, and 6. In total, 844 proteins were identified (≥ 2 specific peptides and FDR < 1% at the protein level).

Quantified protein intensities spanned five orders of magnitude across the 30 samples collected at three time points (Figure 3).

**Figure 3.**
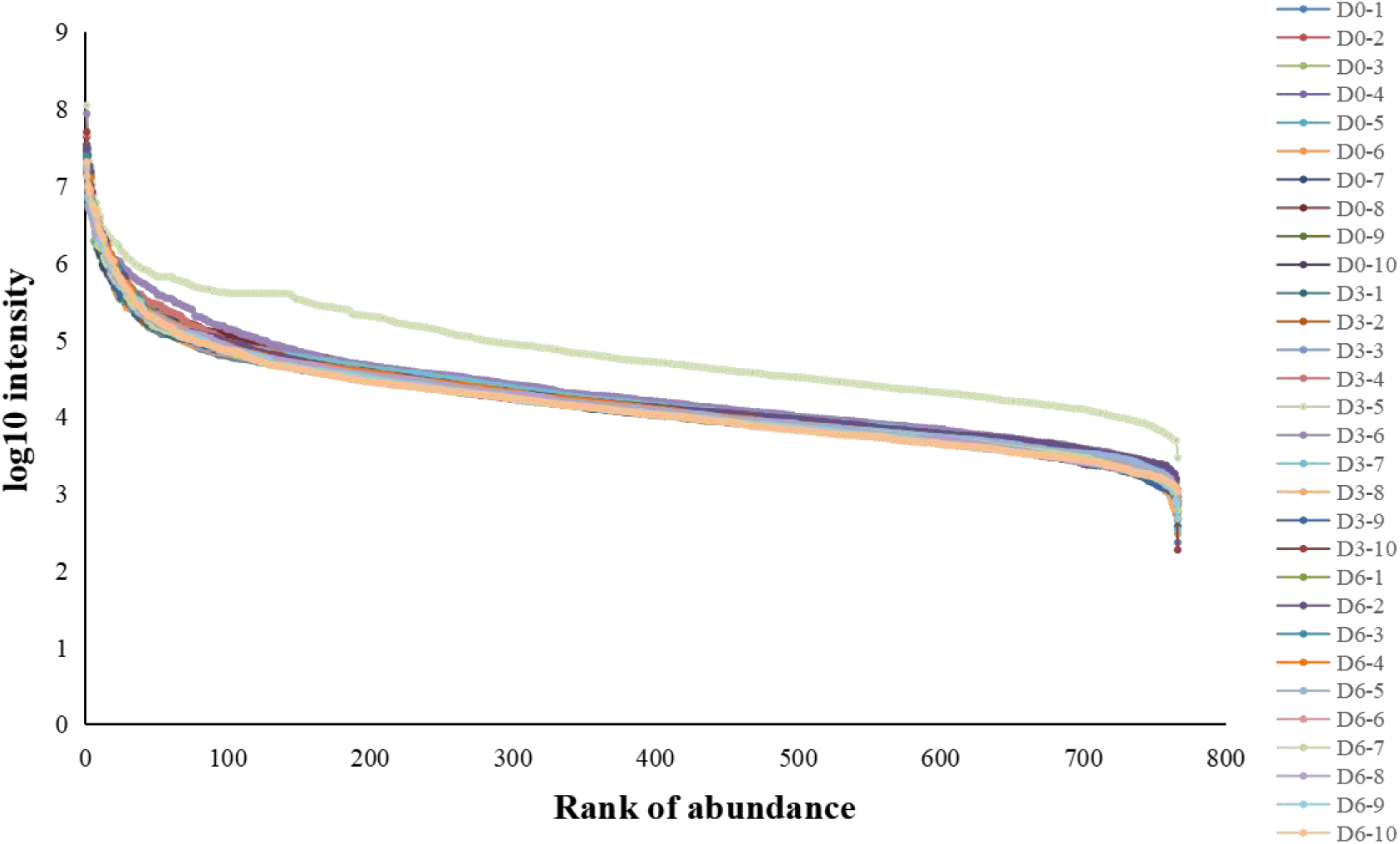
Dynamic ranges of the urinary proteome were measured in urine samples from 10 rats at three time points. Proteins identified in the cohort at all stages were ranked according to their MS signals, which covered more than five orders of magnitude.

#### 3.2.2 Differential proteins at different time points

The urine proteins of rats at different time points of growth and development were differential, and the criteria for screening differential proteins were FC ≥ 1.5 or ≤ 0.67 between groups, two-tailed unpaired t test P < 0.05. The results showed that by comparing Day 0 with Day 3, 37 differential proteins could be identified (Table 1). By comparing Day 3 with Day 6, 75 differential proteins were identified (Table S1). Comparing Day 0 with Day 6, 195 differential proteins were identified (Table S2).

**Table 1.**
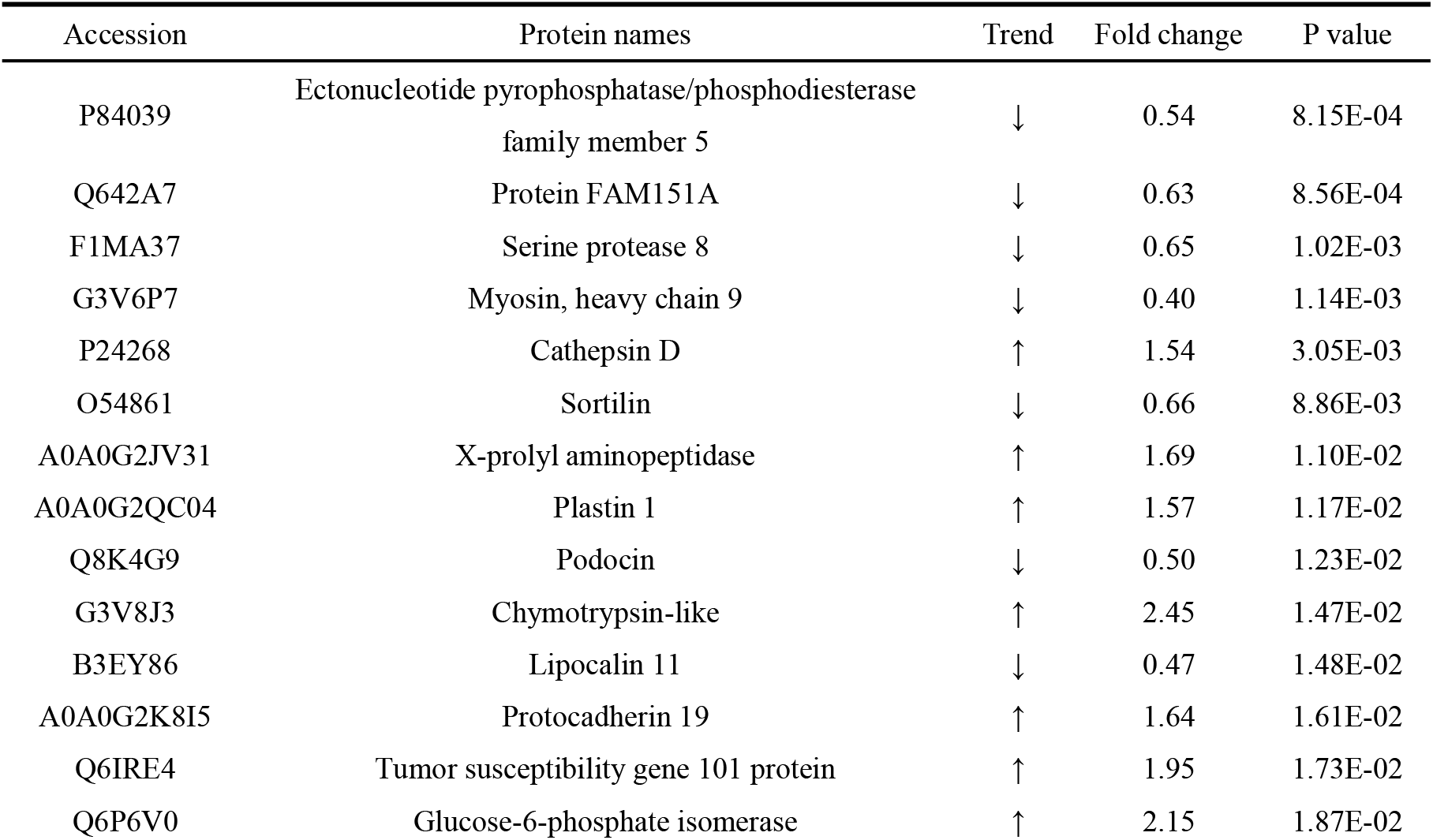
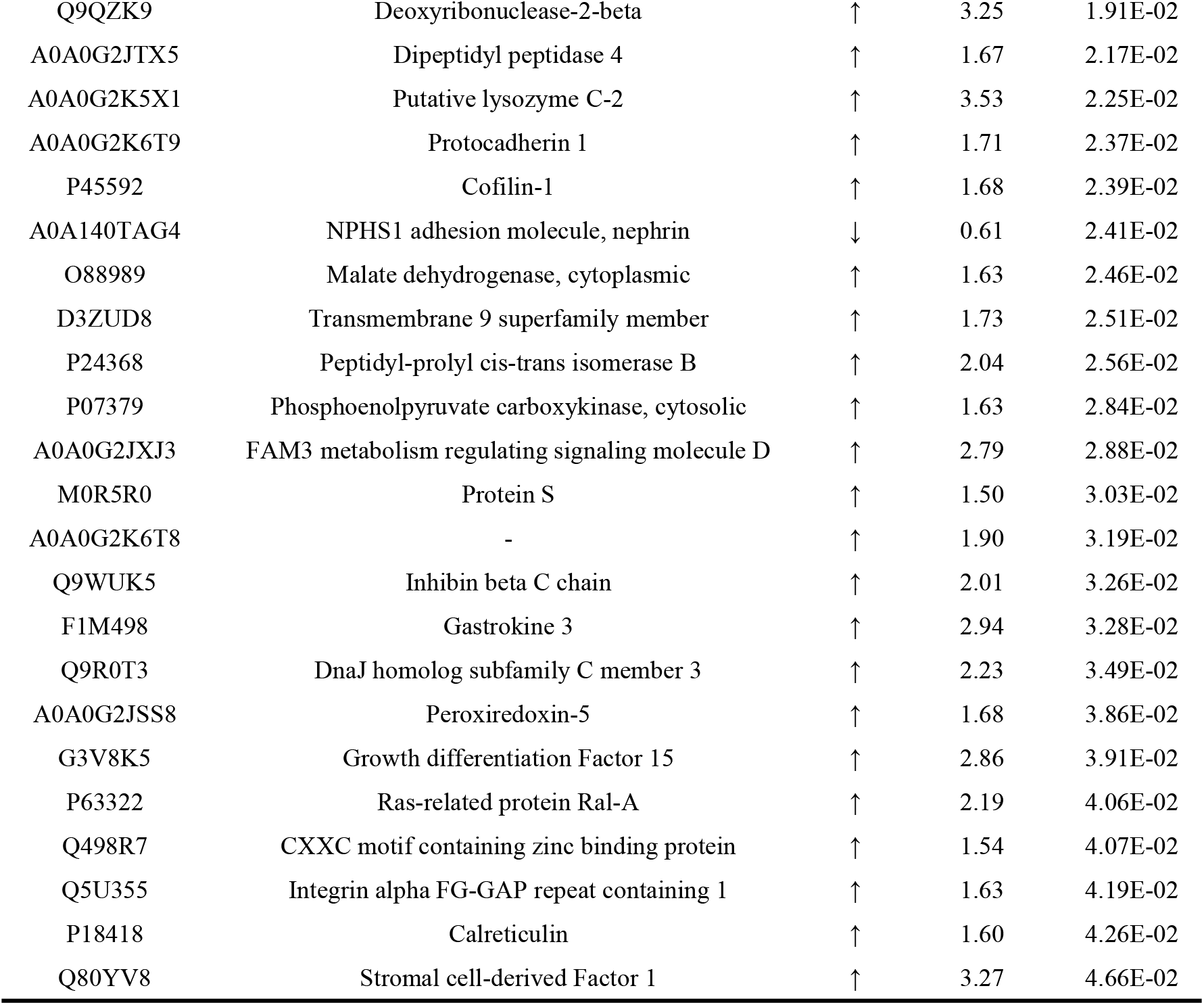
Day 0 vs. Day 3 differential Protein Information (FC≥1.5 或≤0.67, P<0.05)

To further investigate the importance of differential proteins screened at different time points, the screening conditions were changed to FC ≥ 2 or ≤ 0.5, P < 0.01. The results showed that 1 differential protein could be identified between Day 0 and Day 3, 13 differential proteins could be identified between Day 3 and Day 6, and 46 differential proteins could be identified between Day 0 and Day 6 (Table 2). Among them, the latter two time point comparisons presented eight common differential proteins, and their expression trends were completely consistent. By using the UniProt website, five of the downregulated proteins were found to be associated with aging, including broad substrate specificity ATP-binding cassette transporter ABCG2, glutathione hydrolase 1 proenzyme, glutathione synthetase, alpha-crystallin B chain, and glutamate-cysteine ligase regulatory subunit. Eleven differential proteins were found to be associated with developmental pathways, in which Crk-like protein is upregulated and is involved in morphogenesis of various organs in animals as an acetylcholine receptor signaling factor (UniProt website).

**Table 2.**
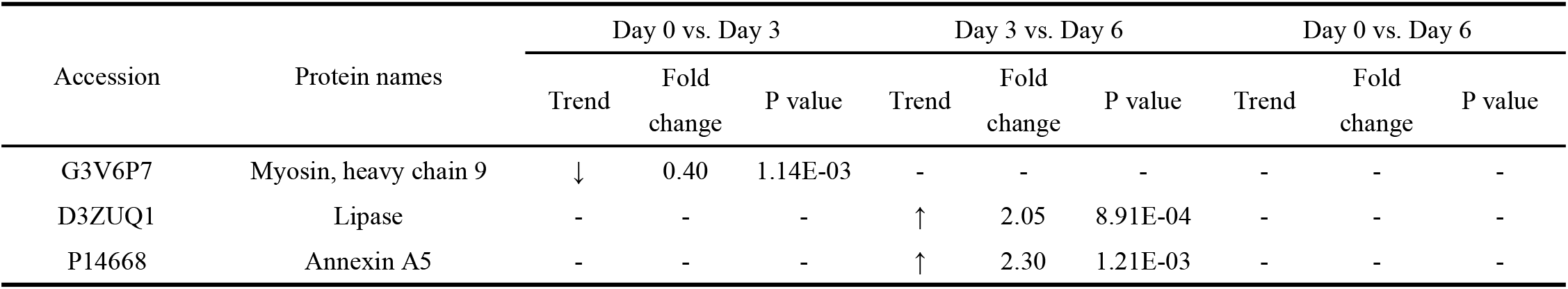
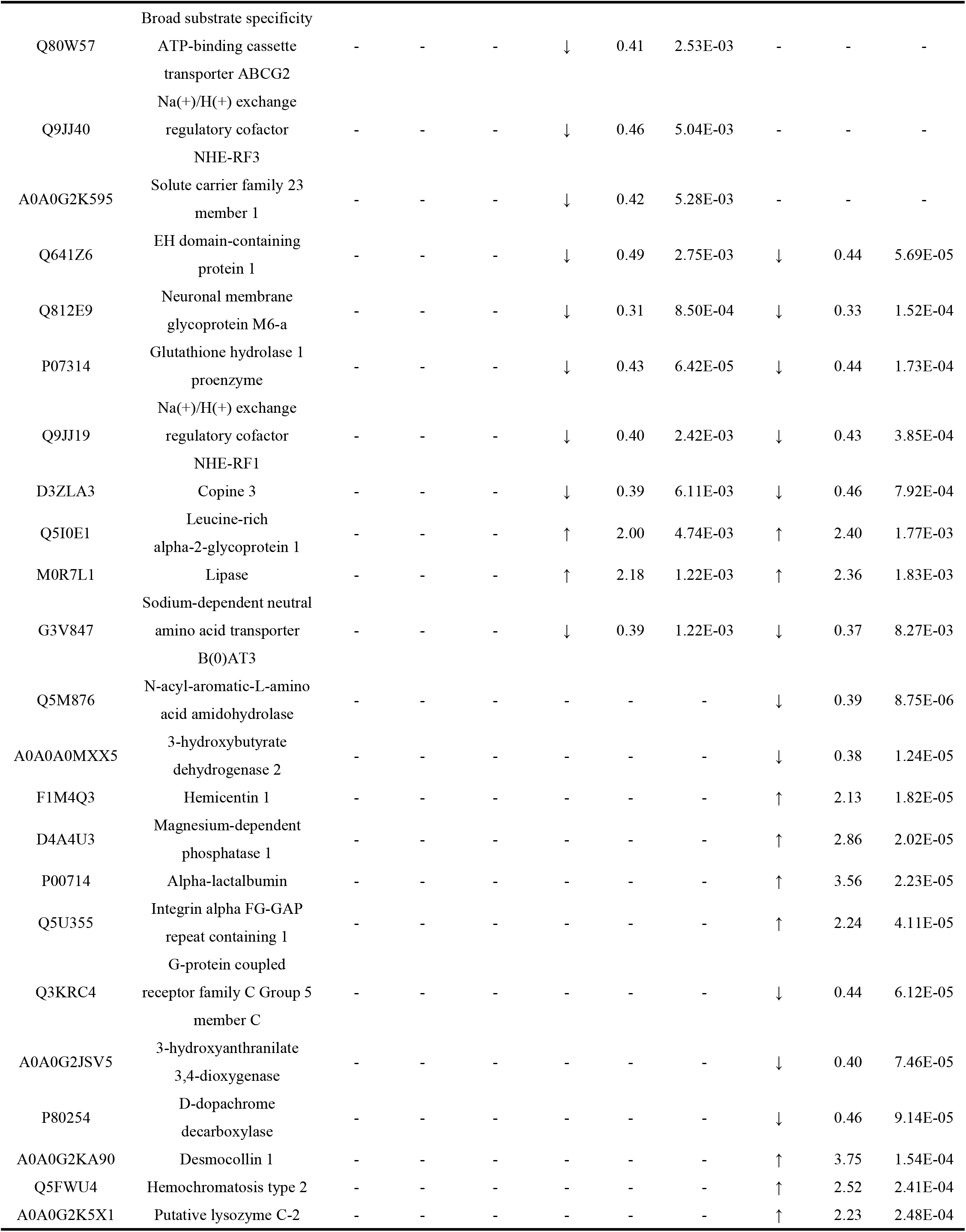
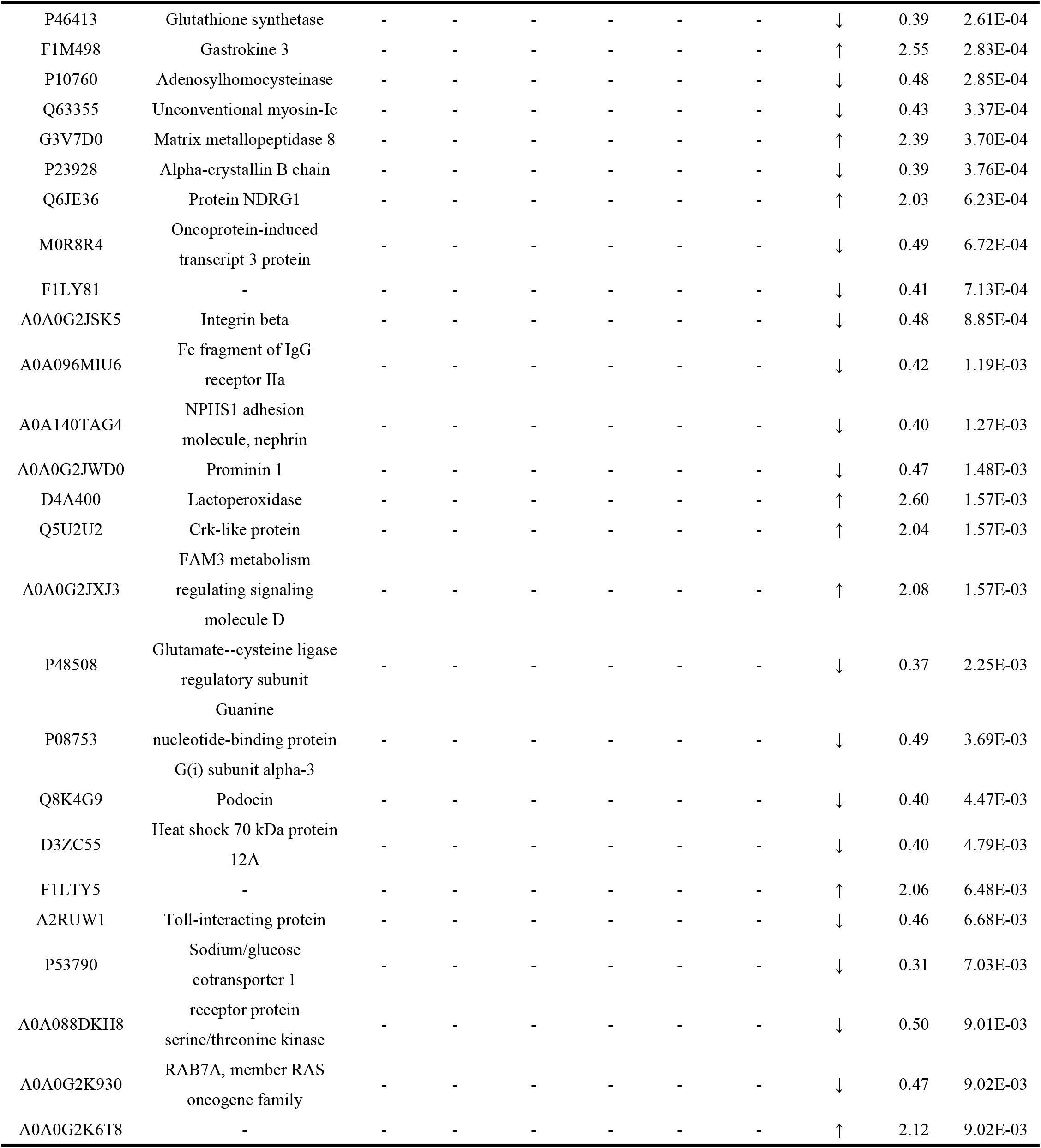
Differential protein information generated by comparison at different time points (FC≥2 或≤0.5, P<0.01)

#### 3.2.3 Randomization test

To determine the possibility that the identified differential proteins were randomly generated, the total proteins identified from 10 samples collected on Day 0 and from 10 samples collected on Day 3 were randomly validated (FC ≥ 1.5 or ≤ 0.67, P < 0.05), and a total of 92,378 different combinations yielded an average of 12.33 differential proteins and 33.32% randomly identified proteins, indicating that at least 66.68% of the differential proteins were not randomly generated (Table 3). Random grouping validation was performed using the total proteins identified from 10 samples collected on Day 3 and from 10 samples collected on Day 6 (FC ≥ 1.5 or ≤ 0.67, P < 0.05), resulting in an average of 13.30 differentially identified proteins and 17.73% randomly identified proteins, indicating that at least 82.27% of differentially identified proteins were not randomly generated (Table 3). Next, random grouping validation was performed under stricter conditions (FC ≥ 2 or ≤ 0.5, P < 0.01), resulting in an average of 0.79 differentially identified proteins and 6.08% randomly identified proteins, indicating that at least 93.92% of differentially identified proteins were not randomly generated (Table 3). Randomization validation was performed on the total proteins identified from 10 samples collected on Day 0 and from 10 samples collected on Day 6 (FC ≥ 1.5 or ≤ 0.67, P < 0.05), yielding an average of 17.99 differentially identified proteins and a proportion of 9.23% randomly identified proteins, indicating that at least 90.77% of differentially identified proteins were not randomly generated (Table 3). Next, randomization validation was performed under more stringent conditions (FC ≥ 2 or ≤ 0.5, P < 0.01), yielding an average of 0.88 differentially identified proteins and a proportion of 1.91% randomly identified proteins, indicating that at least 98.09% of differentially identified proteins were not randomly generated (Table 3).

**Table 3.**
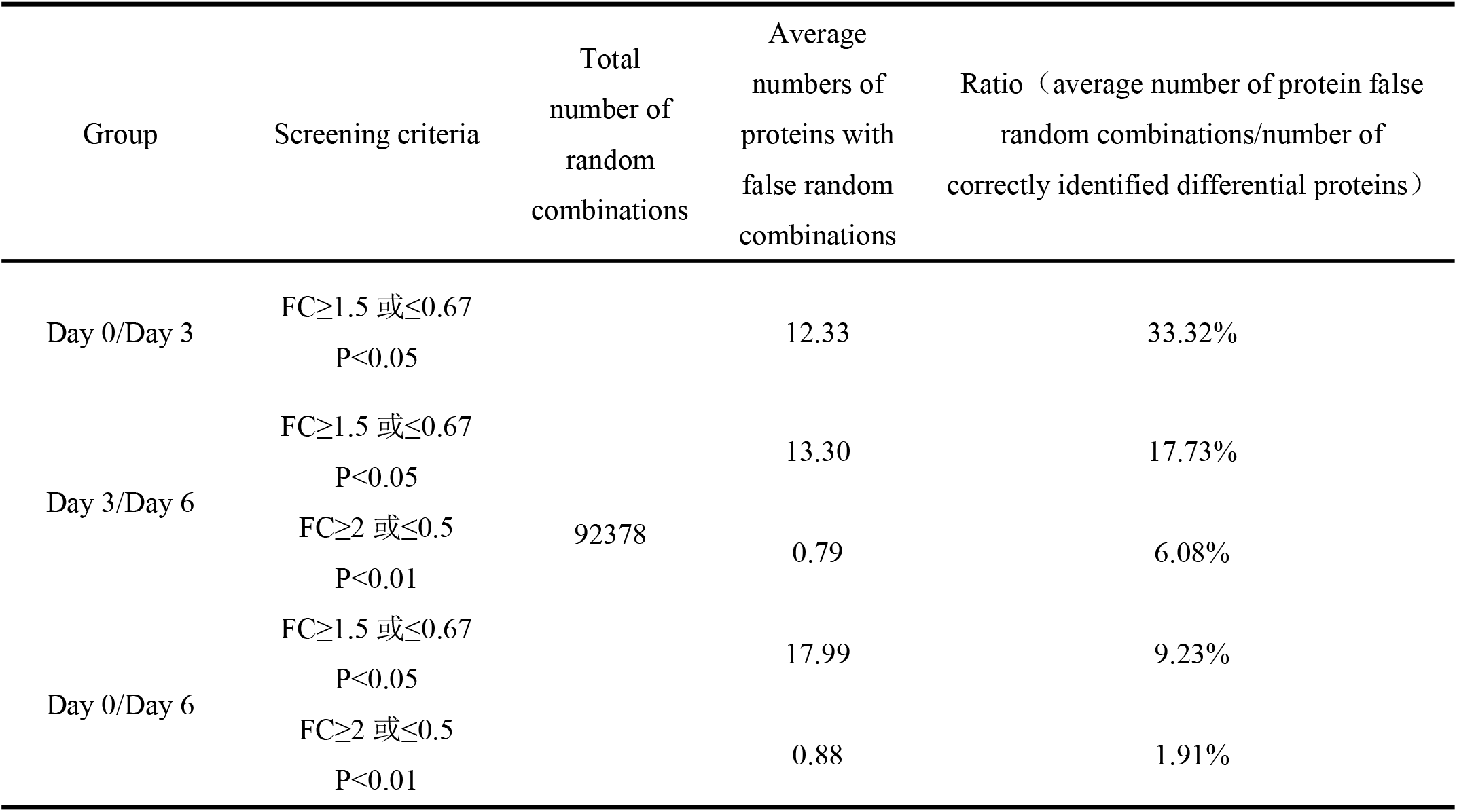
Randomization Results

#### 3.2.4 Functional analysis of differential proteins screened relative to the control at different time points

We used the STRING website (https://cn.string-db.org/), and a PPI (protein-protein interaction) analysis was performed on the differential proteins obtained from the relative ratio screening of Day 0 to Day 6 (FC ≥ 1.5 or ≤ 0.67, P < 0.05) (Figure 4).

**Figure 4.**
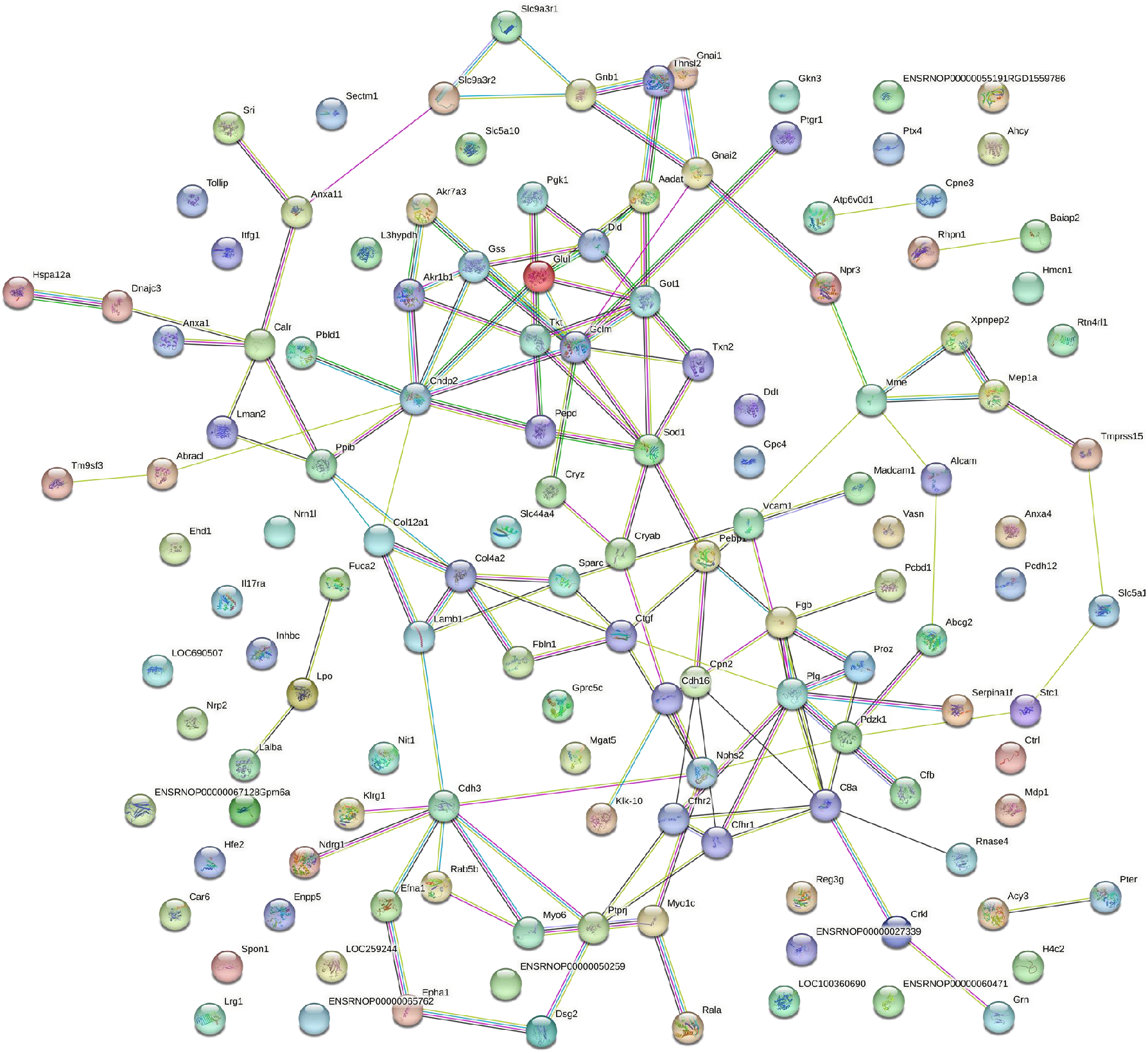
Day 0 vs. Day 6 differential proteins STRING PPI Network Analysis (FC≥1.5 或≤0.67, P<0.05).

To determine the biological relevance of the identified protein networks, we used the OmicsBean website (Fig. http://www.omicsbean.cn/). GO (Gene Ontology) analysis and KEGG pathway analysis were performed on the differential proteins in each group. Some of the strongly enriched biological processes, cellular components, molecular functions, and biological pathways are shown in Figure 5.

**Figure 5.**
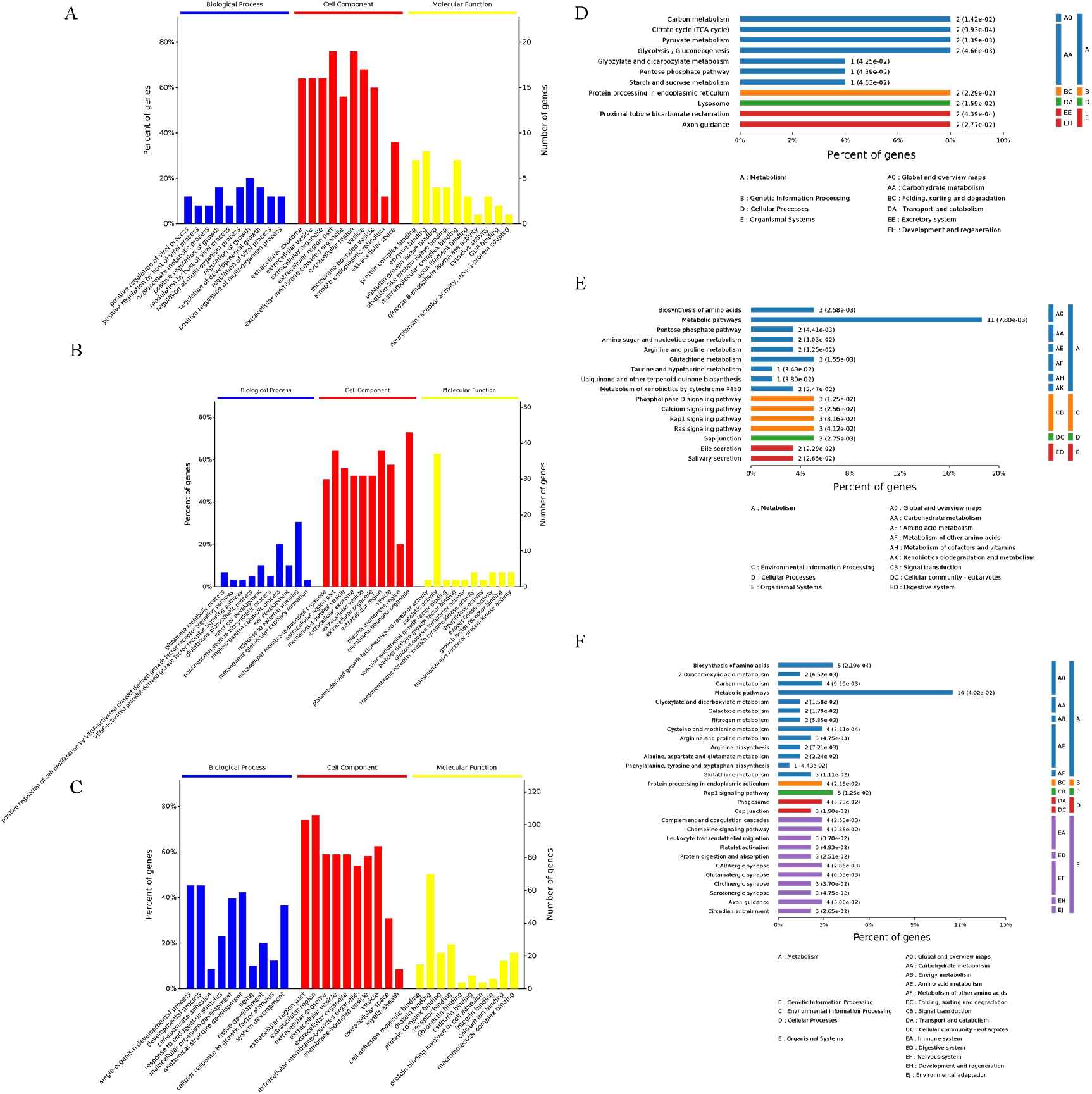
Functional analysis of differential urinary proteins at different time points of rat growth and development. (A) GO analysis of Day 0 vs. Day 3. (B) GO analysis of Day 3 vs. Day 6. (C) GO analysis of Day 0 vs. Day 6. (D) Pathway analysis of Day 0 vs. Day 3. (E) Pathway analysis of Day 3 vs. Day 6. (F) Pathway analysis of Day 0 vs. Day 6.

Differential proteins resulting from the comparison of Day 0 to Day 3 were mainly enriched in biological processes such as regulation of growth, positive regulation of growth, regulation of development growth, regulation of multiorganism processes, and positive regulation of multiorganism processes. Differential proteins resulting from the comparison of Day 3 to Day 6 were mainly enriched in biological processes such as inner ear development, ear development, glutamate metabolic process, and positive regulation of cell proliferation by the VEGF-activated platelet-derived growth factor receptor signaling pathway. Differential proteins resulting from the comparison of Day 0 to Day 6 were enriched in biological processes such as single-organism development process, developmental process, multicellular organism development, anatomical structure development, aging, tissue development, and system development, and several biological processes involved up to 40% of the differential proteins, demonstrating that the reliability of related biological processes was high. In terms of cellular components, the results at the three time points were highly repetitive, and the enriched cellular components mainly included extracellular exosome, extracellular vesicle, extracellular organelle, extracellular region part, extracellular membrane-bound organelle, vesicle, and membrane-bound vesicle. The enriched molecular functions mainly included protein binding, catalytic activity, protein complex binding and macromolecular complex binding. Among the enriched representative signaling pathways, the Rap1 signaling pathway, axon guidance, protein processing in the endoplasmic reticulum, gap junctions, glutathione metabolism, and several other metabolic pathways emerged repeatedly in different time groups.

## 4. Discussion

In this study, mass spectrometry-based nonlabeled quantitative proteomics combined with urine proteomics was used to analyze the rapid growth and development of 10 Wistar rats aged 6-8 weeks for 6 days. Comparing the urine protein levels of rats on Day 0 and Day 3, 37 differential proteins were identified under relaxed screening conditions (FC ≥ 1.5 or ≤ 0.67, P < 0.05) and verified by randomization, with a reliability of 66.68%, while 1 differential protein could be identified under strict screening conditions (FC ≥ 2 or ≤ 0.5, P < 0.01). Comparing Day 3 with Day 6, 75 differential proteins could be identified under relaxed screening conditions (FC ≥ 1.5 or ≤ 0.67, P < 0.05), and the reliability reached 82.27% after randomization, while 13 differential proteins could be identified under strict screening conditions (FC ≥ 2 or ≤ 0.5, P < 0.01). After random grouping, the reliability reached 93.92%. Contrasting Day 0 with Day 6, 195 differential proteins could be identified under relaxed screening conditions (FC ≥ 1.5 or ≤ 0.67, P < 0.05), which were verified by randomization with 90.77% reliability, while 46 differential proteins could be identified under strict screening conditions (FC ≥ 2 or ≤ 0.5, P < 0.01). After random grouping, the reliability reached 98.09%. These results showed that the differential proteins between Day 0, Day 3 and Day 6 were reliably contrasted with each other, the protein reliability also showed an increasing trend with increasing time span, and the differential proteins reliability obtained under strict screening conditions was higher than that obtained under relaxed conditions. The differential proteins we screened excluded the possibility of random production and therefore we inferred that these differential proteins were indeed associated with short-term growth and development in rats. Under stringent screening conditions (FC ≥ 2 or ≤ 0.5, P < 0.01), eight differential proteins were found both in the Day 3 and 6 urine samples. Among them, neuronal membrane glycoprotein M6-a is reported to be a stress-related gene and plays an important role in synapse and filopodia formation, which is essential for development, immunity, angiogenesis, wound healing and metastasis^[14]^. Researchers have found that leucine-rich alpha-2-glycoprotein 1 is able to participate in the regulatory mechanisms of angiogenesis, epithelial-mesenchymal transition (EMT), and apoptosis through the TGF-β (transforming growth factor-β) signaling pathway, affecting tumor development^[15]^. Comparing Day 0 with Day 6, we found that d-3-hydroxybutyrate dehydrogenase 2 was differentially expressed in rat urine under strict screening conditions (FC ≥ 2 or ≤ 0.5, P < 0.01). It has been shown that the regulation of mitochondrial d-3-hydroxybutyrate dehydrogenase can modulate different stages of rat development and physiology^[16]^. Moreover, alpha-lactalbumin can improve energy balance and metabolism^[17]^ and promote the growth and maturation of the small intestine in suckling rats^[18]^. It has also been shown that adenosylhomocysteinase is a biomarker of physical fragility and may play a regulatory role in aging and age-related diseases^[19]^. In addition, NDRG1 (N-myc downstream-regulated gene 1 protein) is an intracellular protein that can be induced under a variety of stress and cell growth regulatory conditions, and NDRG1 is upregulated by cell differentiation signals and inhibits tumor metastasis in various cancer cell lines^[20]^. Win PW et al. showed that β1 integrin beta signaling is required to maintain islet mass and angiogenesis during specific transition windows in developing β-cells^[21]^, while Masuzaki R et al. investigated integrin β1 as a key determinant of liver structure and played a key role as a modulator of TGF-β secretion^[22]^. A review by Barzegar Behrooz A et al. reveals that CD133 (prominin-1) may increase angiogenesis and promote the growth and differentiation of neural cells by activating the Wnt signaling pathway^[23]^. These differences in urinary protein levels may play a key role in the growth and development of rats and explains the sensitivity of urine proteomics, reflects the dynamic changes in the growth and development of rats in just 6 days, and rules out the possibility of random production.

During aging, we identified proteins that were involved in important biological processes, such as growth regulation, positive regulation of growth, development growth regulation, regulation of multiple biological processes, positive regulation of multiple biological processes, inner ear development, ear development, glutamate metabolism, positive regulation of cell proliferation by VEGF-activated platelet-derived growth factor receptor signaling pathway, single biological development process, developmental process, multicellular biological development, anatomical structure development, aging, tissue development, and phylogeny, all of which have some correlation with growth and development. In addition, we found that the RAP1 signaling pathway was enriched. Chrzanowska-Wodnicka M et al.^[24]^ showed that the small GTPase Rap1 is involved in many basic cellular functions and is essential for the development and formation of functional blood vessels in mice. Enriched signaling pathways that were common at multiple time points include axonal guidance, protein processing in the endoplasmic reticulum, gap junctions, glutathione metabolism, and several other metabolic pathways.

These results suggest that the urine proteome can reflect the dynamic changes in rats during normal growth and development in a short period of time. We believe that the rate of body changes may be different at different stages of rat development, and the results of this study on male Wistar rats with rapid growth and development at 6-8 weeks of age provide a means for in-depth and meticulous study of growth and development. It also suggests a key interference factor for future animal experiments using urinary protein biomarkers and questions if there is a suitable time period of rat growth and development and the aging period that most greatly reduces the interference of rat growth and development and aging on the experiment.

## 5. Conclusion

This study showed that the urine proteome can reflect the growth- and development-related dynamic changes in rats in a rather short period of time. Therefore, the effect of short-term development on urinary protein needs to be taken into account when performing marker studies using model animals that grow rapidly.

## Authorship Contribution Statement

**Yuqing Liu:** Conceptualization, Investigation, Writing-original draft, Data curation, Visualization. **Minhui Yang:** Investigation. **Haitong Wang:** Investigation. **Yuzhen Chen:** Investigation. **Youhe Gao:** Conceptualization, Writing-review & editing, Supervision, Funding acquisition.

## Acknowledgments

This work was supported by the National Key Research and Development Program of China (2018YFC0910202), the Fundamental Research Funds for the Central Universities (2020KJZX002), and the Beijing Natural Science Foundation (7172076).

---

**Table S1.** Day 3 vs. Day 6 differential protein information (FC≥1.5 或≤0.67, P<0.05)

| Accession | Protein names | Trend | Fold change | P value |
| --- | --- | --- | --- | --- |
| P07314 | Glutathione hydrolase 1 proenzyme | ↓ | 0.43 | 6.42E-05 |
| Q63041 | Alpha-1-macroglobulin | ↑ | 1.71 | 2.89E-04 |
| Q812E9 | Neuronal membrane glycoprotein M6-a | ↓ | 0.31 | 8.50E-04 |
| D3ZUQ1 | Lipase | ↑ | 2.05 | 8.91E-04 |
| P14668 | Annexin A5 | ↑ | 2.30 | 1.21E-03 |
| G3V847 | Sodium-dependent neutral amino acid transporter B(0)AT3 | ↓ | 0.39 | 1.22E-03 |
| M0R7L1 | Lipase | ↑ | 2.18 | 1.22E-03 |
| A0A0A0MXT4 | Solute carrier organic anion transporter family member | ↓ | 0.53 | 1.95E-03 |
| A0A0G2JSJ2 | Cytidine/uridine monophosphate kinase 1 | ↓ | 0.62 | 2.12E-03 |
| A0A0G2KB26 | Matrix remodeling-associated protein 8 | ↑ | 1.53 | 2.40E-03 |
| Q9JJ19 | Na(+)/H(+) exchange regulatory cofactor NHE-RF1 | ↓ | 0.40 | 2.42E-03 |
| Q80W57 | Broad substrate specificity ATP-binding cassette transporter ABCG2 | ↓ | 0.41 | 2.53E-03 |
| Q641Z6 | EH domain-containing protein 1 | ↓ | 0.49 | 2.75E-03 |
| D4AA31 | Prolylcarboxypeptidase | ↑ | 1.66 | 3.81E-03 |
| D3ZE16 | Alpha-L-iduronidase | ↓ | 0.66 | 4.13E-03 |
| F7F389 | Complement C9 | ↑ | 1.82 | 4.22E-03 |
| Q5M876 | N-acyl-aromatic-L-amino acid amidohydrolase | ↓ | 0.52 | 4.71E-03 |
| Q5I0E1 | Leucine-rich alpha-2-glycoprotein 1 | ↑ | 2.00 | 4.74E-03 |
| Q9JJ40 | Na(+)/H(+) exchange regulatory cofactor NHE-RF3 | ↓ | 0.46 | 5.04E-03 |
| A0A0G2K595 | Solute carrier family 23 member 1 | ↓ | 0.42 | 5.28E-03 |
| D3ZLA3 | Copine 3 | ↓ | 0.39 | 6.11E-03 |
| G3V7D0 | Matrix metalloproteinase 8 | ↑ | 1.69 | 6.91E-03 |
| A0A140TAG4 | NPHS1 adhesion molecule, nephrin | ↓ | 0.66 | 7.13E-03 |
| A0A0G2QC04 | Plastin 1 | ↓ | 0.61 | 7.25E-03 |
| D3ZH39 | receptor protein-tyrosine kinase | ↑ | 1.57 | 9.17E-03 |
| D4A400 | Lactoperoxidase | ↑ | 1.96 | 9.70E-03 |
| Q5M8C3 | Serine | ↓ | 0.63 | 1.04E-02 |
| A0A0G2JZ18 | Pentraxin 4 | ↑ | 1.58 | 1.11E-02 |
| F1MAD3 | Polycystin 1, transient receptor potential channel interacting | ↓ | 0.64 | 1.13E-02 |
| P61459 | Pterin-4-alpha-carbinolamine dehydratase | ↓ | 0.53 | 1.15E-02 |
| G3V6A0 | Platelet-derived growth factor receptor alpha | ↓ | 0.50 | 1.18E-02 |
| Q6Q0N1 | Cytosolic nonspecific dipeptidase | ↓ | 0.66 | 1.23E-02 |
| P48508 | Glutamate--cysteine ligase regulatory subunit | ↓ | 0.50 | 1.25E-02 |
| P53790 | Sodium/glucose cotransporter 1 | ↓ | 0.39 | 1.37E-02 |
| G3V8X5 | Solute carrier family 5 | ↓ | 0.47 | 1.39E-02 |
| A0A0G2JWD0 | Prominin 1 | ↓ | 0.48 | 1.47E-02 |
| P80254 | D-dopachrome decarboxylase | ↓ | 0.55 | 1.55E-02 |
| A0A0A0MXX5 | 3-hydroxybutyrate dehydrogenase 2 | ↓ | 0.45 | 1.60E-02 |
| F1M978 | Inositol-1-monophosphatase | ↓ | 0.66 | 1.69E-02 |
| A0A0G2JT43 | Solute carrier family 2, facilitated glucose transporter member 5 | ↓ | 0.47 | 1.74E-02 |
| G3V7L4 | Cadherin 16 | ↓ | 0.61 | 1.86E-02 |
| F1LQI1 | Hydroxyacyl glutathione hydrolase | ↓ | 0.62 | 2.03E-02 |
| P50137 | Transketolase | ↑ | 1.76 | 2.26E-02 |
| P97546 | Neuroplastin | ↓ | 0.57 | 2.39E-02 |
| Q5U362 | Annexin | ↓ | 0.62 | 2.43E-02 |
| A0A0G2JSV5 | 3-hydroxyanthranilate 3,4-dioxygenase | ↓ | 0.51 | 2.47E-02 |
| Q6MG71 | Choline transporter-like protein 4 | ↓ | 0.32 | 2.48E-02 |
| Q9WUW9 | Sulfotransferase 1C2A | ↓ | 0.50 | 2.55E-02 |
| P09606 | Glutamine synthetase | ↓ | 0.47 | 2.69E-02 |
| P04904 | Glutathione S-transferase alpha-3 | ↓ | 0.65 | 2.75E-02 |
| D4A4U3 | Magnesium-dependent phosphatase 1 | ↑ | 1.68 | 2.92E-02 |
| F1LVR0 | IgLON family member 5 | ↓ | 0.40 | 3.13E-02 |
| P01681 | Ig kappa chain V region S211 | ↑ | 1.93 | 3.36E-02 |
| E9PU23 | Delta-like protein | ↓ | 0.47 | 3.45E-02 |
| B0BNJ1 | LOC683667 protein | ↓ | 0.36 | 3.45E-02 |
| D3ZV91 | trans-L-3-hydroxyproline dehydratase | ↓ | 0.45 | 3.61E-02 |
| G3V6D9 | Na(+)/H(+) exchange regulatory cofactor NHE-RF | ↓ | 0.51 | 3.61E-02 |
| A1L1J8 | RAB5B, member RAS oncogene family | ↓ | 0.48 | 3.64E-02 |
| Q64602 | Kynurenine/alpha-aminoadipate aminotransferase, mitochondrial | ↓ | 0.55 | 3.66E-02 |
| P51607 | N-acylglucosamine 2-epimerase | ↓ | 0.42 | 3.76E-02 |
| P02770 | Albumin | ↓ | 0.53 | 3.80E-02 |
| D3ZXY4 | Aldehyde dehydrogenase 8 family, member A1 | ↓ | 0.53 | 3.81E-02 |
| Q6AXM6 | Intercellular adhesion molecule 2 | ↓ | 0.65 | 3.84E-02 |
| A0A0G2JSQ7 | Kallikrein 1-related peptidase B3 | ↓ | 0.55 | 3.86E-02 |
| P38918 | Aflatoxin B1 aldehyde reductase member 3 | ↓ | 0.63 | 3.93E-02 |
| Q3KRC4 | G-protein coupled receptor family C Group 5 member C | ↓ | 0.60 | 4.10E-02 |
| Q5FVI6 | V-type proton ATPase subunit C 1 | ↓ | 0.44 | 4.10E-02 |
| Q4V7D9 | Acid sphingomyelinase-like phosphodiesterase | ↓ | 0.62 | 4.24E-02 |
| Q499Q4 | Phosphoglucomutase 1 | ↓ | 0.46 | 4.37E-02 |
| Q05030 | Platelet-derived growth factor receptor beta | ↓ | 0.66 | 4.40E-02 |
| D4A8C5 | 1-phosphatidylinositol 4,5-bisphosphate phosphodiesterase | ↓ | 0.39 | 4.40E-02 |
| Q5I0D7 | Xaa-Pro dipeptidase | ↓ | 0.23 | 4.41E-02 |
| G3V8L3 | Lamin A, isoform CRA_b | ↓ | 0.38 | 4.64E-02 |
| P16975 | SPARC | ↓ | 0.51 | 4.72E-02 |
| P32755 | 4-hydroxyphenylpyruvate dioxygenase | ↓ | 0.57 | 4.85E-02 |

**Table S2.**
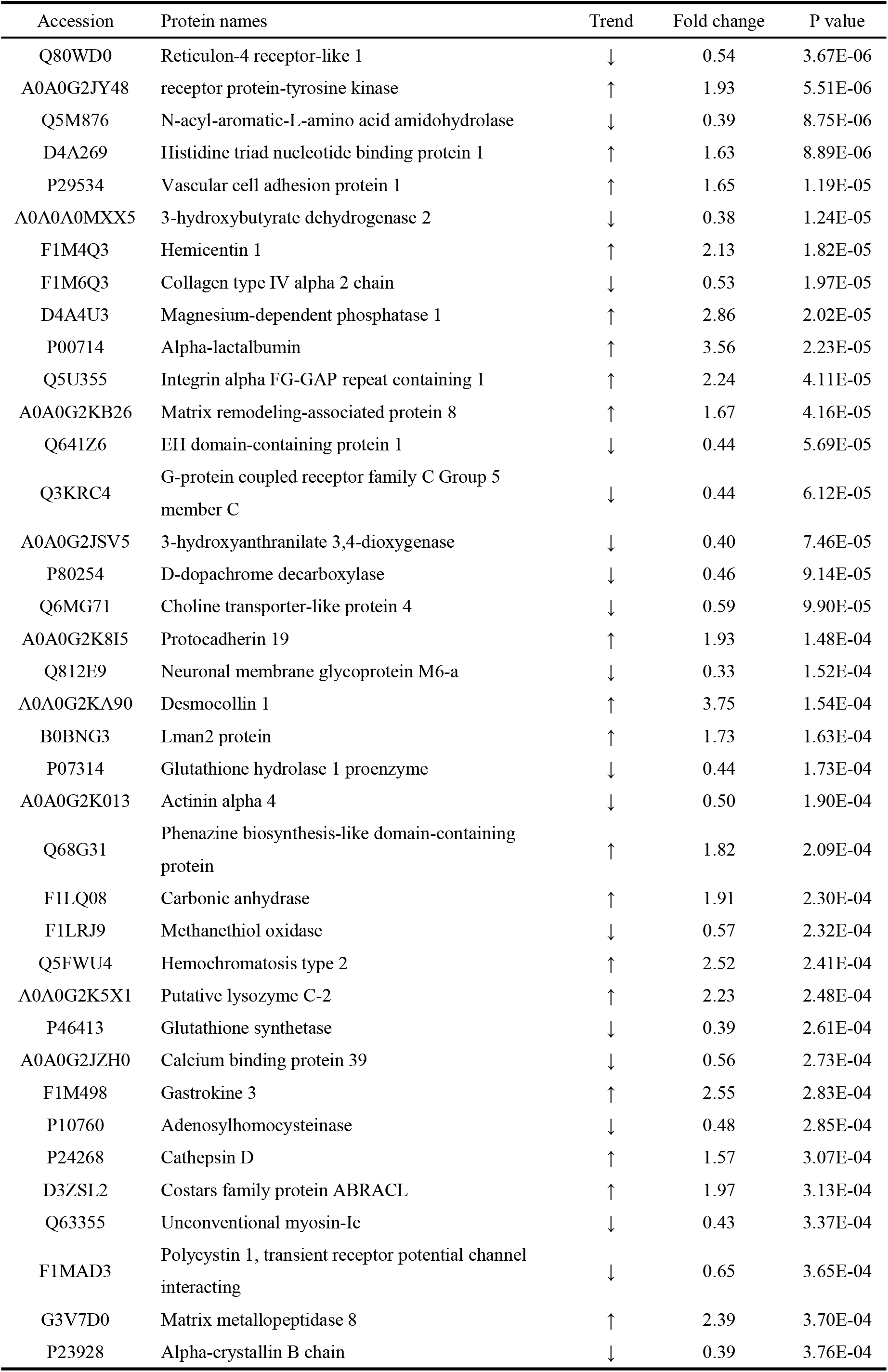
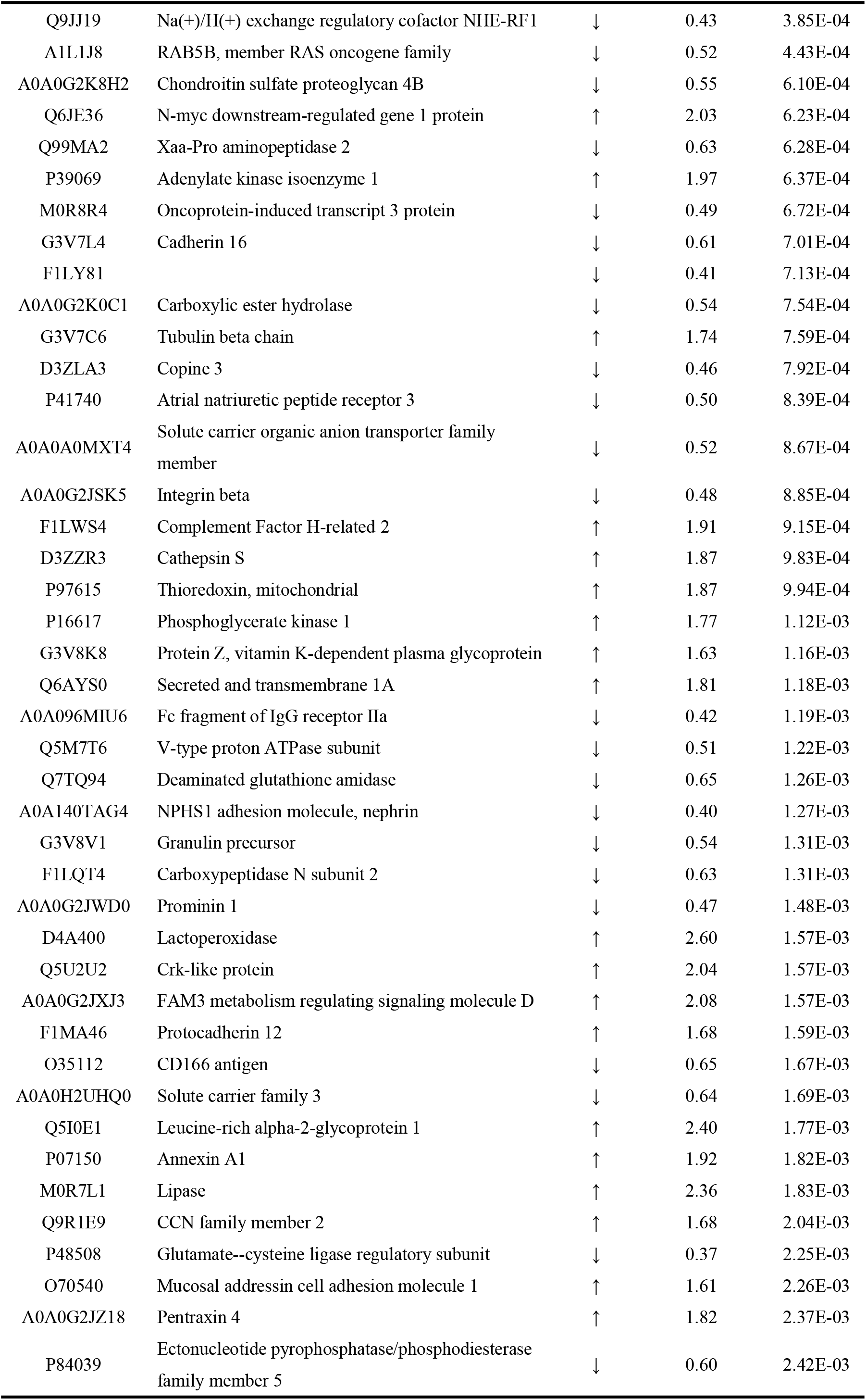
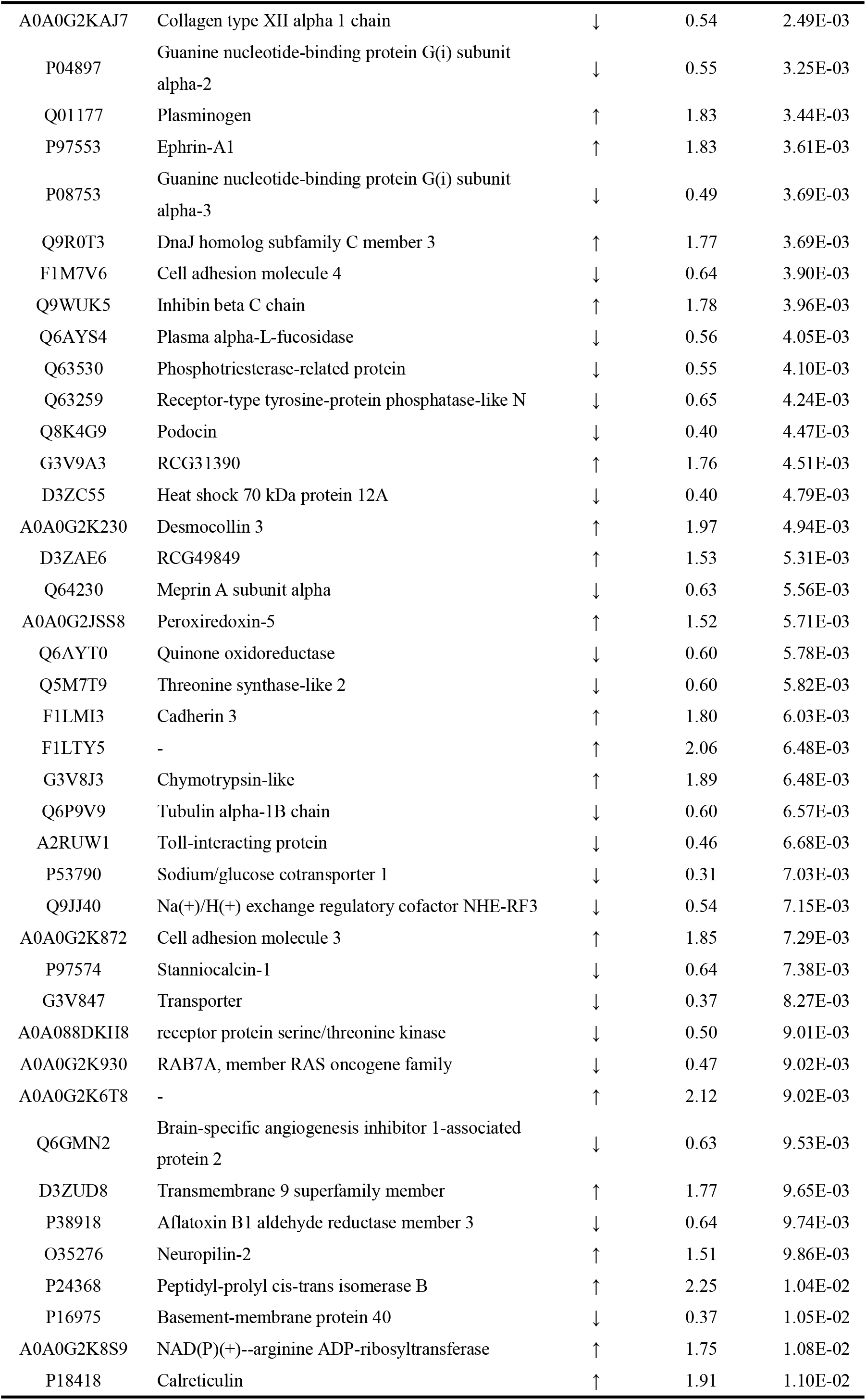
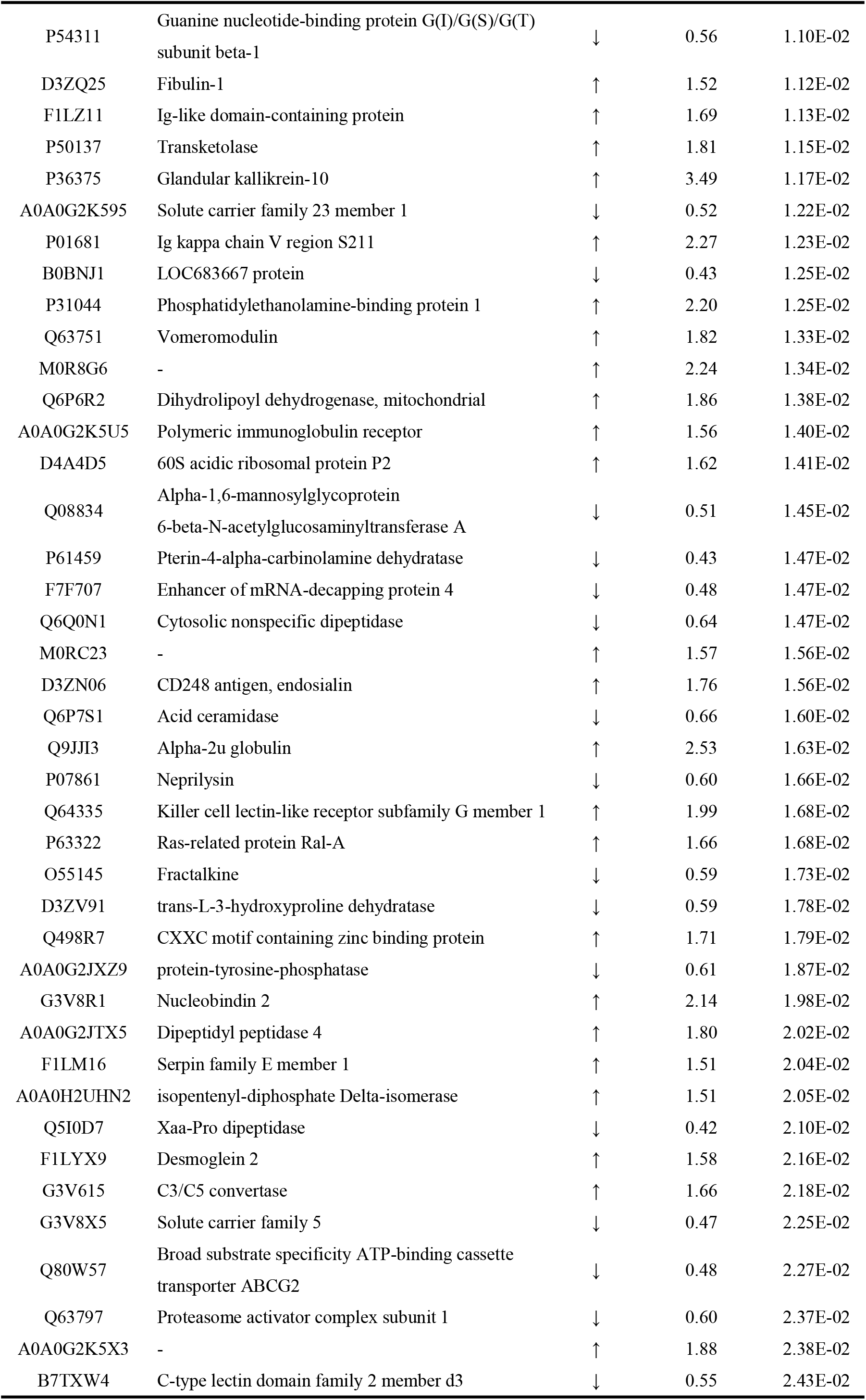
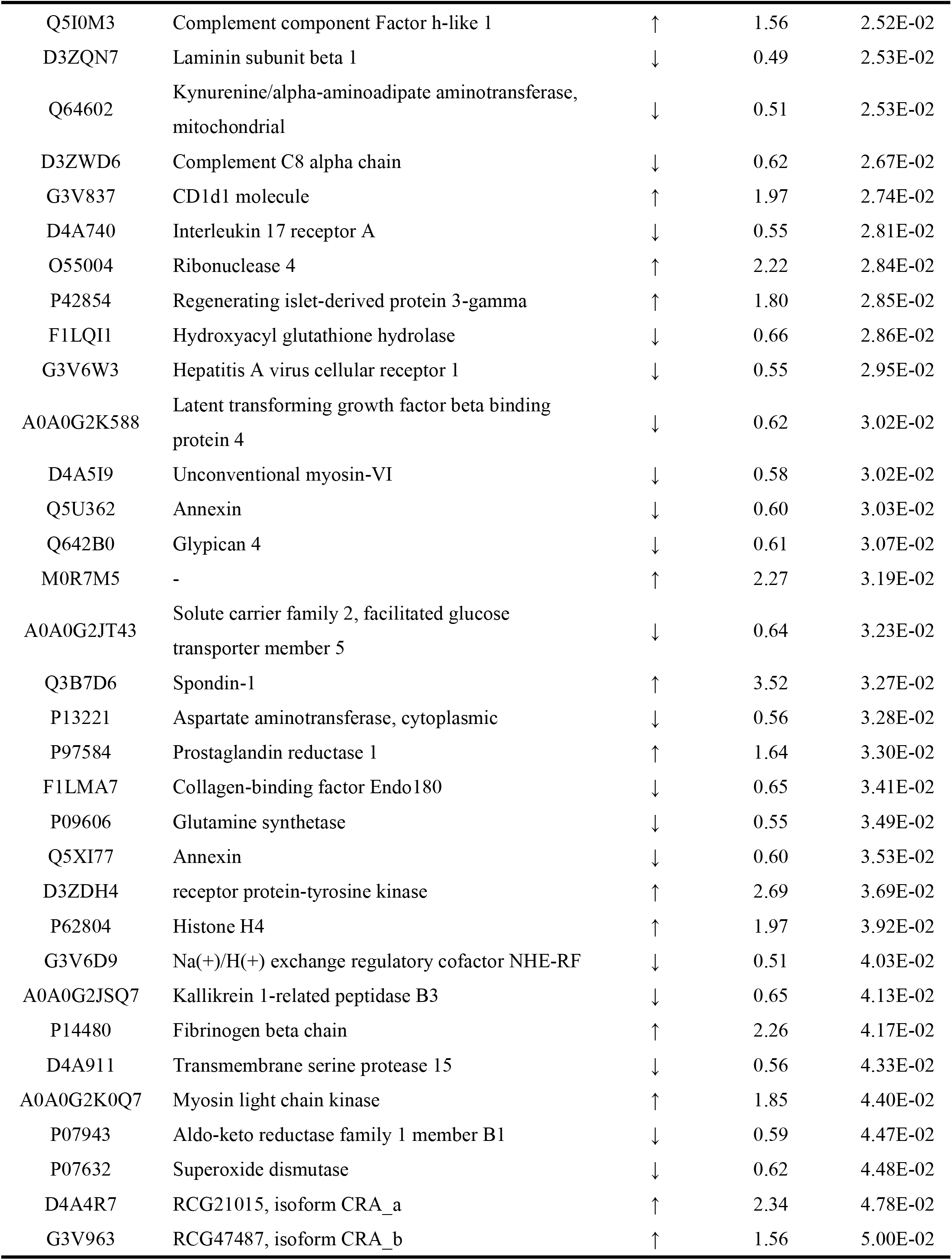
Day 0 vs. Day 6 differential protein information (FC≥1.5 或≤0.67, P<0.05)

